# Sex differences in central and peripheral fatigue induced by sustained isometric ankle plantar flexion

**DOI:** 10.1101/2021.09.03.458912

**Authors:** Donguk Jo, Miriam Goubran, Martin Bilodeau

**Author notes:** Corresponding author Donguk Jo.

## Abstract

The main aim of this study was to determine sex differences in central and peripheral fatigue produced by a sustained isometric exercise of ankle plantar flexors in healthy young adults. Ten males and fourteen females performed a sustained isometric ankle exercise until task failure. Maximal voluntary isometric contraction torque (plantarflexion), voluntary activation level (using the twitch interpolation technique), and twitch contractile properties (twitch peak torque, twitch half relaxation time, and low frequency fatigue index) were measured before, immediately after, and throughout a recovery period (1, 2, 5, and 10 min) following the exercise protocol in order to characterize neuromuscular fatigue. Fatigue had a significant effect (p ≤ 0.05) on all dependent variables. Other than for the maximal voluntary contraction torque, where males showed a greater fatigue-related decrease than females, males and females showed generally similar changes with fatigue. Altogether, our findings indicate no major differences in central or peripheral fatigue mechanisms between males and females to explain a somewhat greater fatigability in males.

## Introduction

The limits of an individual’s performance during athletic events, occupational tasks, daily activities, or exercise can be partly explained by fatigue that leads to acute changes in the neuromuscular system (1). Neuromuscular fatigue can be defined as any exercise-induced loss of ability to generate force or power within a muscle or muscle group (2,3). Neuromuscular fatigue develops progressively from the onset of exercise until an individual may no longer be able to perform a given task. This fatigue progression can result from changes in neurophysiological mechanisms at different levels and sites (e.g., central and peripheral nervous system and within the muscle) before the capacity to produce a given force level significantly decreases (2).

In an attempt to organize and clarify the origins of these changes, neuromuscular fatigue has been separated into two components, central and peripheral fatigue. Central fatigue represents a progressive exercise-induced reduction in voluntary activation (or reduced neural drive to the muscle) (3,4). A decrease in voluntary activation could arise from changes in the activation/commands from the brain to an alteration in motor neuron activity, and may involve afferent feedback from muscles (4). This reduction in voluntary activation is reflected by an increase in the superimposed twitch torque produced by peripheral (motor) nerve stimulation during maximal voluntary isometric contractions (MVICs) of the fatigued muscle or muscle group (5,6). This reduced voluntary activation indicates that motor units are not fully recruited or firing quickly enough to produce optimal (maximal) muscle torque when the stimulation is delivered (7). In contrast, peripheral fatigue is caused by processes occurring at or distal to the neuromuscular junction (3). Peripheral fatigue can be assessed by looking at changes in twitch contractile properties of a given fatigued muscle by electrically stimulating the peripheral nerve while the muscle is at rest. Typical fatigue-related changes include: reduced twitch peak torque (8), increased twitch half relaxation time (the time to have half of the decrease in a twitch peak torque) (9,10) and decreased low frequency fatigue (LFF) index (ratio of low frequency twitch torque to high frequency twitch torque) (11).

Sex differences in neuromuscular fatigue have been found for some fatiguing tasks, although the differences vary depending on muscle group, exercise intensity, and contraction type (12). Typically, males show greater fatigability (greater reduction in muscle force) than females, not only for isometric but also dynamic fatiguing tasks such as: single muscle group isometric contraction (e.g., elbow flexion, ankle dorsiflexion) (13,14) and slow-to-moderate velocity contractions (e.g., knee flexion/extension, elbow flexion) (15,16). The greater fatigability in males compared with females has been suggested to be mainly due to a greater oxidative metabolic capacity (17), a greater proportion of fatigue-resistant muscle fibers (type I) within skeletal muscles (12), and lesser voluntary activation during fatiguing tasks (14) in females compared with males.

Amongst the limited number of studies that have determined sex differences in central and peripheral fatigue, inconsistent results can be found with regards to the contribution of central versus peripheral fatigue mechanisms, even when observing lesser fatigue resistance (e.g., greater exercise-induced decreases in MVIC torque or shorter exercise time to failure) in males than females (12,18). Some studies using isometric exercise protocols of the lower limb (14,19), for example, found greater central fatigue (greater reductions in voluntary activation) following the exercise protocol in males compared with females, but showed no significant sex differences in peripheral fatigue (similar reductions in resting twitch peak torque in both sex groups). In contrast, several studies using isometric exercise protocols of the upper/lower limb (20–24) found greater exercise-induced peripheral fatigue in males compared with females, reporting greater reductions in resting twitch peak torque and slowing of twitch relaxation following exercise in males compared with females, while not observing sex differences in central fatigue.

These contrasting findings in a limited number of studies highlight the need for further evidence with regards to sex differences in neuromuscular fatigue mechanisms in healthy adults. In particular, sex differences in neuromuscular fatigue have not been studied extensively for ankle plantar flexors (12). Given the primary role of ankle plantar flexors in standing stability and gait performance (25,26), specific knowledge concerning neuromuscular fatigue of this muscle group can contribute to inform optimal strategies for exercise/rehabilitation in both males and females. Therefore, our main aim was to determine sex differences in central and peripheral fatigue of ankle plantar flexors during a sustained isometric exercise in healthy young adults.

## Methods

### Participants

Ten healthy young males (27.3 ± 5.3 years; 183.5 ± 5.8 cm; 91.5 ± 11.4 kg) and fourteen healthy young females (29.4 ± 4.9 years; 167.4 ± 7.3 cm; 62.9 ± 9.8 kg) volunteered for this study. Participants had no neurological disorders or musculoskeletal injuries to the lower leg, as confirmed by a health questionnaire. These participants showed moderate-to-high physical activity levels (8.63 ±2.17 out of 15), as reported on a habitual physical activity questionnaire (27). The study was approved by the University of Ottawa and Bruyère Continuing Care research ethics boards. Participants were informed of all procedures before participation in the study.

### Experimental procedure

All participants volunteered for a single session. The experimental protocol began by determining the supramaximal intensity of electrical stimulation at the popliteal fossa needed to generate the maximal twitch torque from ankle plantar flexors of dominant leg (see below for details). Participants then familiarized themselves with the main task by generating bilateral submaximal intermittent isometric contractions of ankle plantar flexors in a sitting position on a dynamometer. Once comfortable with the task, they were asked to perform three MVICs in ankle plantar flexion (approximately 3-5 s in duration), with a 2-min interval between each MVIC. During each MVIC, one doublet stimulus was delivered at the maximum (plateau) of the MVIC torque to record a superimposed doublet torque (the dominant leg). Immediately after each MVIC, one doublet stimulus and then one singlet stimulus were delivered with a 2-s interval while the ankle plantar flexors were at rest in order to measure resting singlet/doublet twitch peak torque. After these pre-fatigue measures, participants performed a sustained isometric ankle plantarflexion with both legs in a sitting position until task failure in order to produce plantar flexors fatigue (see below for details). Post-fatigue MVICs were then performed after 1, 2, 5, and 10 min of recovery following the fatiguing exercise. Doublet and singlet electrical stimuli were delivered during and after each MVIC as was done for the pre-fatigue measures, as well as immediately after the exercise (Fig 1).

**Fig 1.**
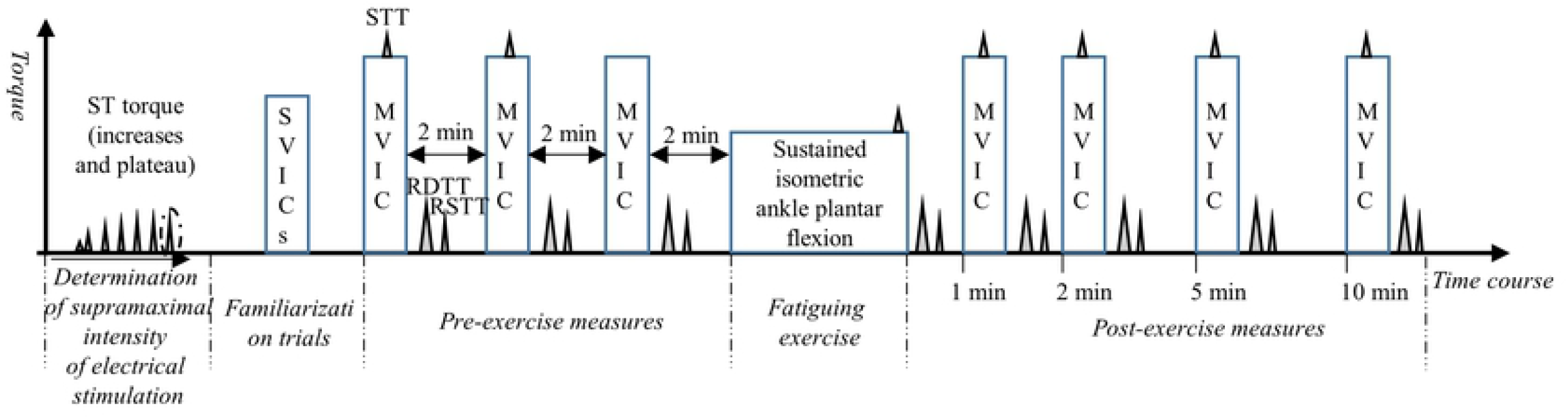
Schematic representation of the experimental procedure. The supramaximal intensity of singlet electrical stimuli generating maximal twitch torque (circled) was set before the pre-fatigue measures. The supramaximal intensity was maintained for all subsequent singlet and doublet stimuli. Submaximal voluntary isometric contractions (SVICs) for familiarization trials were performed. Superimposed twitch torque (STT) induced by doublet stimuli at the maximum (plateau) of maximal voluntary isometric contractions (MVICs), and resting doublet twitch torque (RDTT) and resting singlet twitch torque (RSTT) immediately after MVICs and at the end of the fatiguing exercise were recorded.

### MVIC and twitch torque recording

A dynamometer (Biodex System 3 Pro ®, Biodex medical systems, Shirley, USA) was used to record MVIC peak torque, superimposed twitch torque, and resting doublet and singlet twitch peak torque (all isometric). Participants sat on the dynamometer chair with their legs extended forward and both feet placed on the footplate (aligning the external malleoli with the center of rotation of the leg apparatus). Their feet, chest, and thighs were secured with non-elastic straps.

### Electrical stimulation

Two stimulating electrodes (diameter: 8 mm) were placed longitudinally along the length of the posterior tibial nerve in the popliteal fossa of the dominant leg. A constant current stimulator (Digitimer DS7AH, Digitimer Ltd, Hertfordshire, United Kingdom) delivered single (pulse duration = 200 µs) and doublet electrical pulses (two consecutive singlet stimuli separated by 10 ms). In order to determine a supramaximal intensity before the pre-fatigue measurements, singlet stimulus intensity was increased until an increase in voltage no longer generated an increment in twitch torque amplitude. The voltage was then increased by 10 to 15% to ensure supramaximal intensity, which was maintained for all subsequent stimuli.

### Fatiguing exercise protocol

A sustained isometric exercise protocol was performed to produce fatigue of ankle plantar flexors. Participants were asked to sit on the dynamometer and keep pushing the footplate (plantarflexion) with both feet for as long as possible while producing a target torque corresponding to 30% of MVIC torque represented by a cursor on a computer screen. The fatiguing protocol stopped when participants no longer produced and maintained the target torque for 3 consecutive seconds, at which point a superimposed doublet was provided to estimate voluntary activation at the end of the fatigue task (before the participants were told to relax). Verbal encouragement was given throughout the fatiguing exercise.

### Data analysis

MVIC peak torque, voluntary activation, peak torque of resting singlet and doublet twitches, half relaxation time of singlet and doublet twitch torques, and a low frequency fatigue ratio were obtained before, immediately after the fatiguing exercise (or at the end of the exercise for voluntary activation only), and throughout a 10-min recovery period (1, 2, 5, and 10 min). The half-relaxation time is the time from the twitch peak torque to a 50% reduction in torque amplitude (28). Low frequency fatigue is quantified by a ratio of resting singlet twitch peak torque to resting doublet twitch peak torque (11). Voluntary activation was calculated using the formula (3):

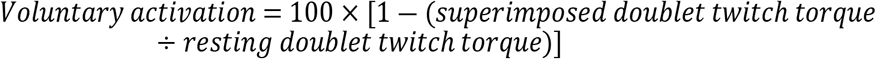

### Statistical analysis

Minimum sample size required was calculated using G* power (version 3.1.9.4) and shown to be at least 10 participants per group, with a power of 0.95, an effect size of 0.71, and alpha of 0.05. The effect size was estimated from existing data, with respect to sex differences in fatigue-induced reductions in voluntary activation (19).

Mixed-model ANOVAs with one between-subjects (sex: males vs. females) and one within-subjects [repeated measures (fatigue): pre-fatigue vs. fatigue (immediately or at the end of the exercise) vs. recovery period (1, 2, 5, and 10 min)] factor, were used to analyze the effect of fatigue, sex, and their interaction on the following dependent variables: MVIC peak torque, resting singlet/doublet twitch peak torque, voluntary activation, low frequency fatigue ratio and half-relaxation time of resting singlet/doublet twitch torque. In cases where Mauchly’s test of Sphericity (within-subjects effect) was significant, the Greenhouse-Geisser correction was used to determine significance. Pairwise comparisons and t-tests (independent) were performed as appropriate/when significance was detected from the ANOVAs using a Bonferroni correction. A Pearson correlation analysis and a multiple linear regression analysis were also performed to assess the association/contribution of changes in voluntary activation (central fatigue) and twitch torque (peripheral fatigue) to changes in MVIC torque after 1 min of recovery. The alpha level chosen for all analyses was set at 0.05.

## Results

### Sex effect

There were some differences between males and females. Males presented with ∼48%, ∼89%, and ∼80% greater values compared with women for MVIC torque, doublet torque and singlet torque respectively (Figs 2-4, Table 1). Doublet and singlet half-relaxation times were significantly longer by 20% (doublet) and about 27% (singlet) in women compared with men (Figs 5 and 6, Table 1). No significant sex effect was found for VA, and LFF ratio (Figs 7 and 8, Table1). The time needed to reach the 70% decrement in MVIC torque (fatigue task duration) was somewhat longer in females (229 ± 76 s) than males (189 ± 64 s). However, this did not reach statistical significance (t = -1.12, p = 0.28).

**Table 1.** Statistical results of the 2-way ANOVAs.

| Variables | Fatigue |  |  | Sex |  |  | Fatigue X Sex |  |  |
| --- | --- | --- | --- | --- | --- | --- | --- | --- | --- |
|  | *F* | *P-value* | *Partial Eta*<sup>2</sup> | *F* | *P-value* | *Partial Eta*<sup>2</sup> | *F* | *P value* | *Partial Eta*<sup>2</sup> |
| MVIC torque | 30.90 | **<0.001** | 0.58 | 12.92 | **0.002** | 0.37 | 4.24 | **0.02** | 0.16 |
| VA | 37.47 | **<0.001** | 0.63 | 1.72 | 0.204 | 0.08 | 0.61 | 0.61 | 0.03 |
| DT | 11.30 | **<0.001** | 0.34 | 37.63 | **<0.001** | 0.63 | 2.38 | 0.07 | 0.10 |
| ST | 21.56 | **<0.001** | 0.50 | 25.10 | **<0.001** | 0.53 | 1.28 | 0.29 | 0.06 |
| LFF ratio | 4.42 | **0.02** | 0.17 | 2.02 | 0.17 | 0.08 | 1.12 | 0.34 | 0.05 |
| DTHRT | 82.98 | **<0.001** | 0.79 | 4.16 | **0.05** | 0.16 | 0.12 | 0.93 | 0.01 |
| STHRT | 46.59 | **<0.001** | 0.68 | 5.26 | **0.03** | 0.19 | 1.01 | 0.39 | 0.04 |

MVIC: maximal voluntary isometric contraction; DT: resting doublet twitch peak torque; ST: resting singlet twitch peak torque; VA: voluntary activation; LFF: low frequency fatigue ratio; DTHRT: doublet twitch half-relaxation time; and STHRT: singlet twitch half-relaxation time.

**Fig 2.**
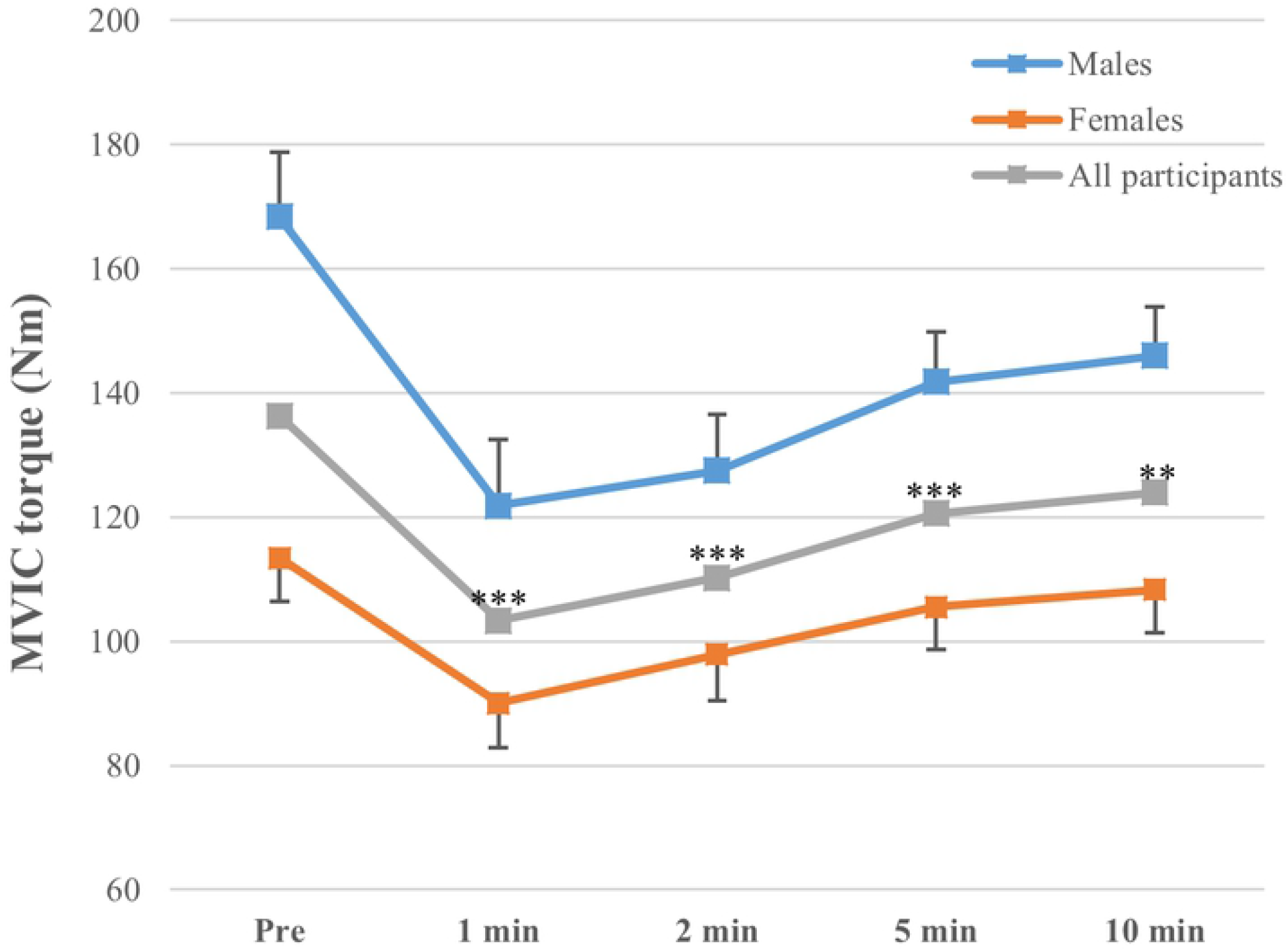
Changes in maximal voluntary isometric contraction (MVIC) peak torque with fatigue and recovery. *** (*p*<0.001) and ** (*p*<0.01) indicate a significant decrease in MVIC peak torque compared with baseline torque (Pre). There was a significant sex X fatigue interaction for MVIC peak torque, with males presenting with a greater decrease in MVC peak torque post-fatigue compared with females. Post hoc tests showed that the change from pre-fatigue to 5 min of recovery was significantly greater in males compared with females (p = 0.012) and approached significance for the change from pre-fatigue to 2 min of recovery (p = 0.06).

**Fig 3.**
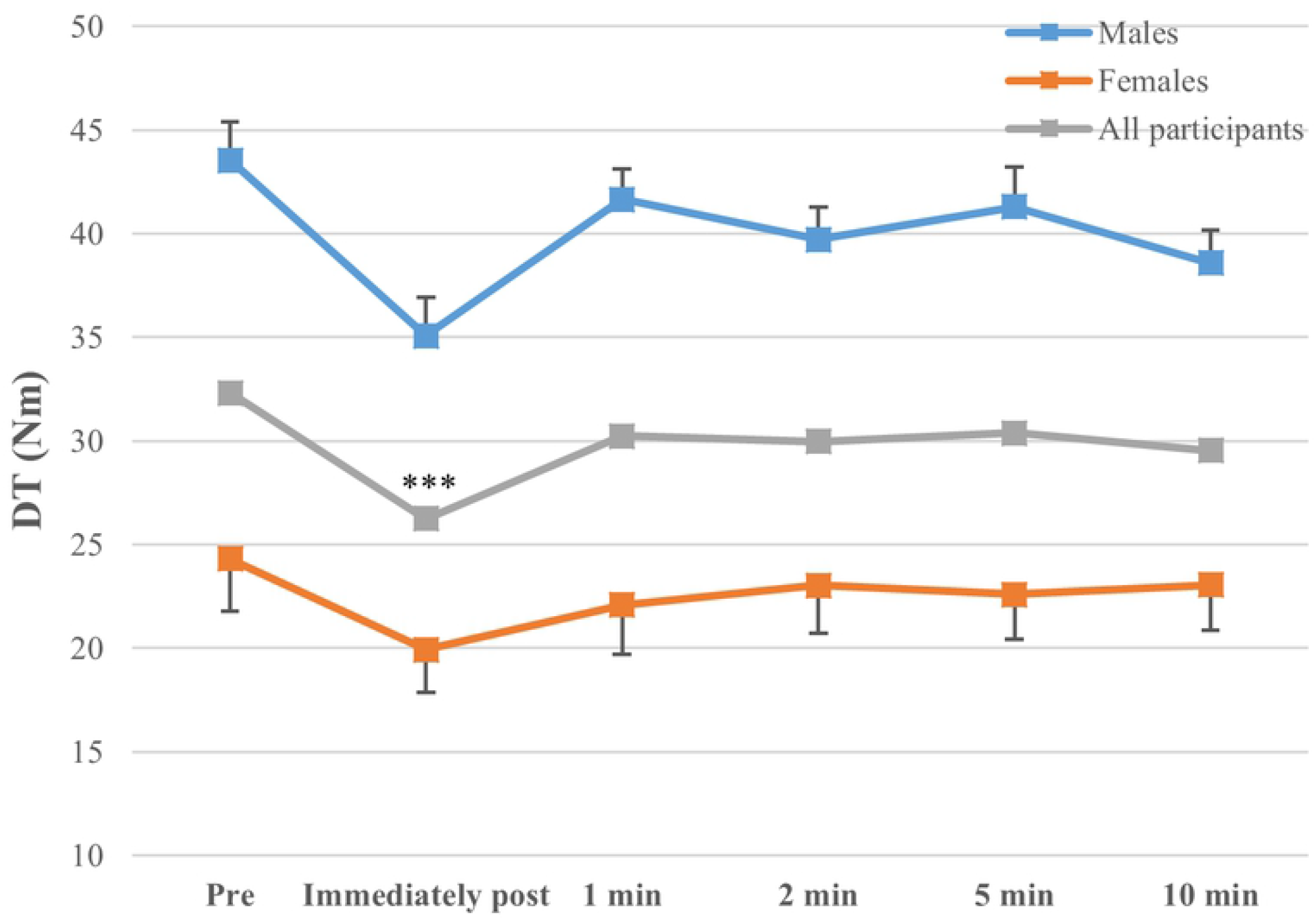
Changes in resting doublet twitch peak torque (DT) with fatigue and recovery. *** (*p*<0.001) indicates a significant decrease in the twitch peak torque post-exercise compared with baseline values (Pre).

**Fig 4.**
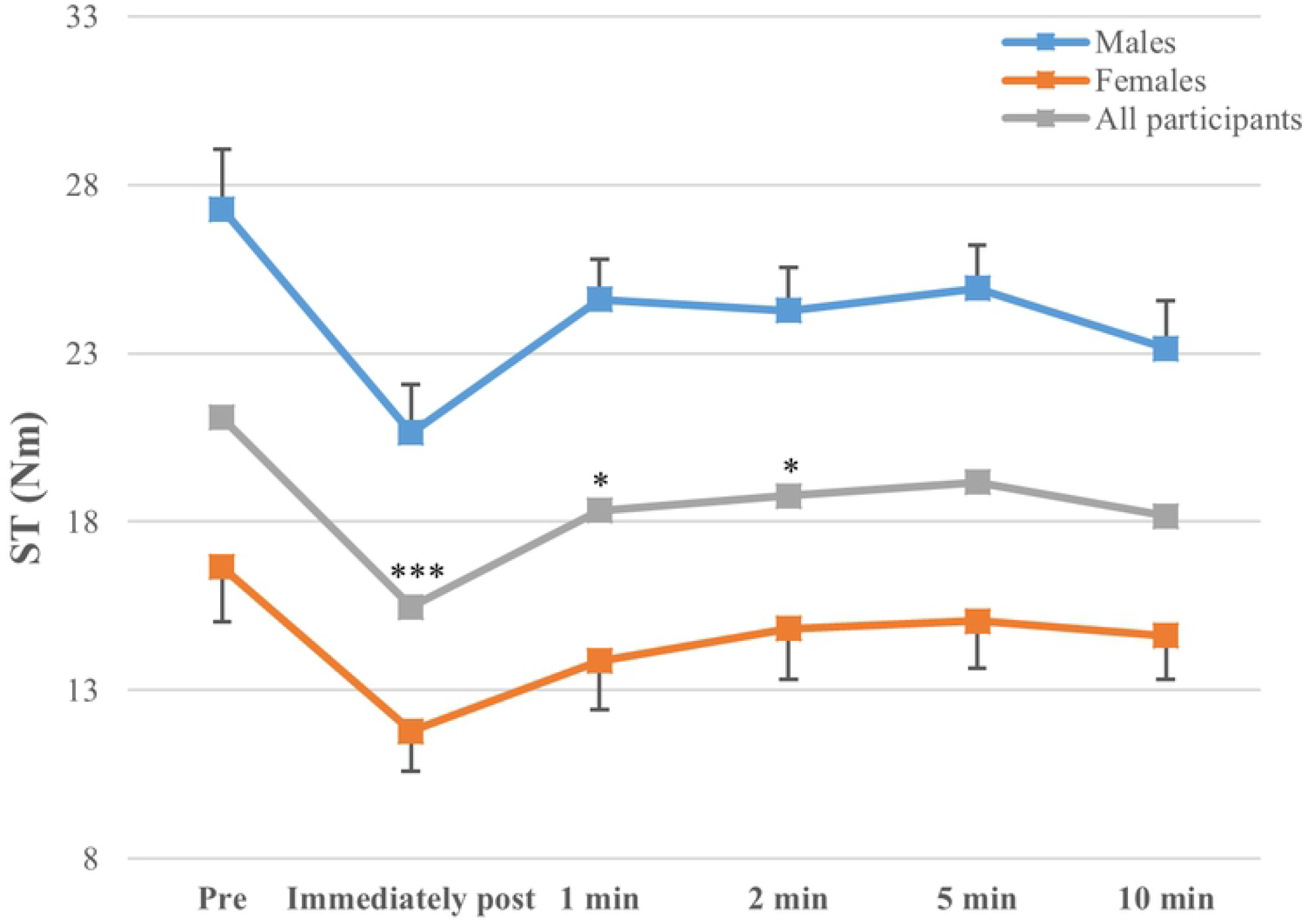
Changes in resting singlet twitch peak torque (ST) with fatigue and recovery. *** (*p*<0.001) and * (*p*<0.05) indicate a significant reduction in the twitch peak torque following exercise compared with baseline values (Pre).

**Fig 5.**
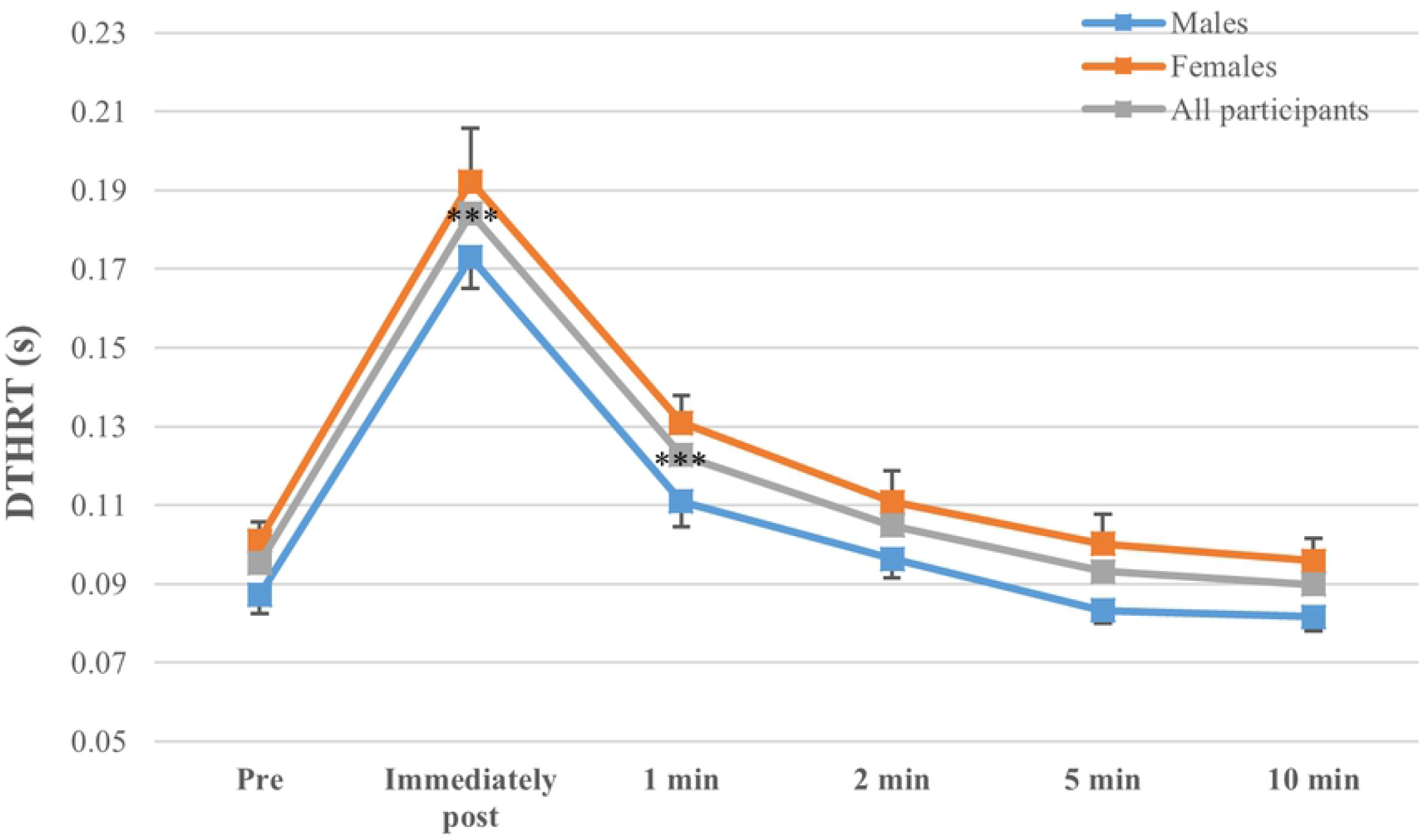
Changes in doublet twitch torque half-relaxation time (DTHRT) with fatigue and recovery. *** (*p*<0.001) indicates a significant difference in the half-relaxation time at post-exercise compared with pre-exercise values (Pre).

**Fig 6.**
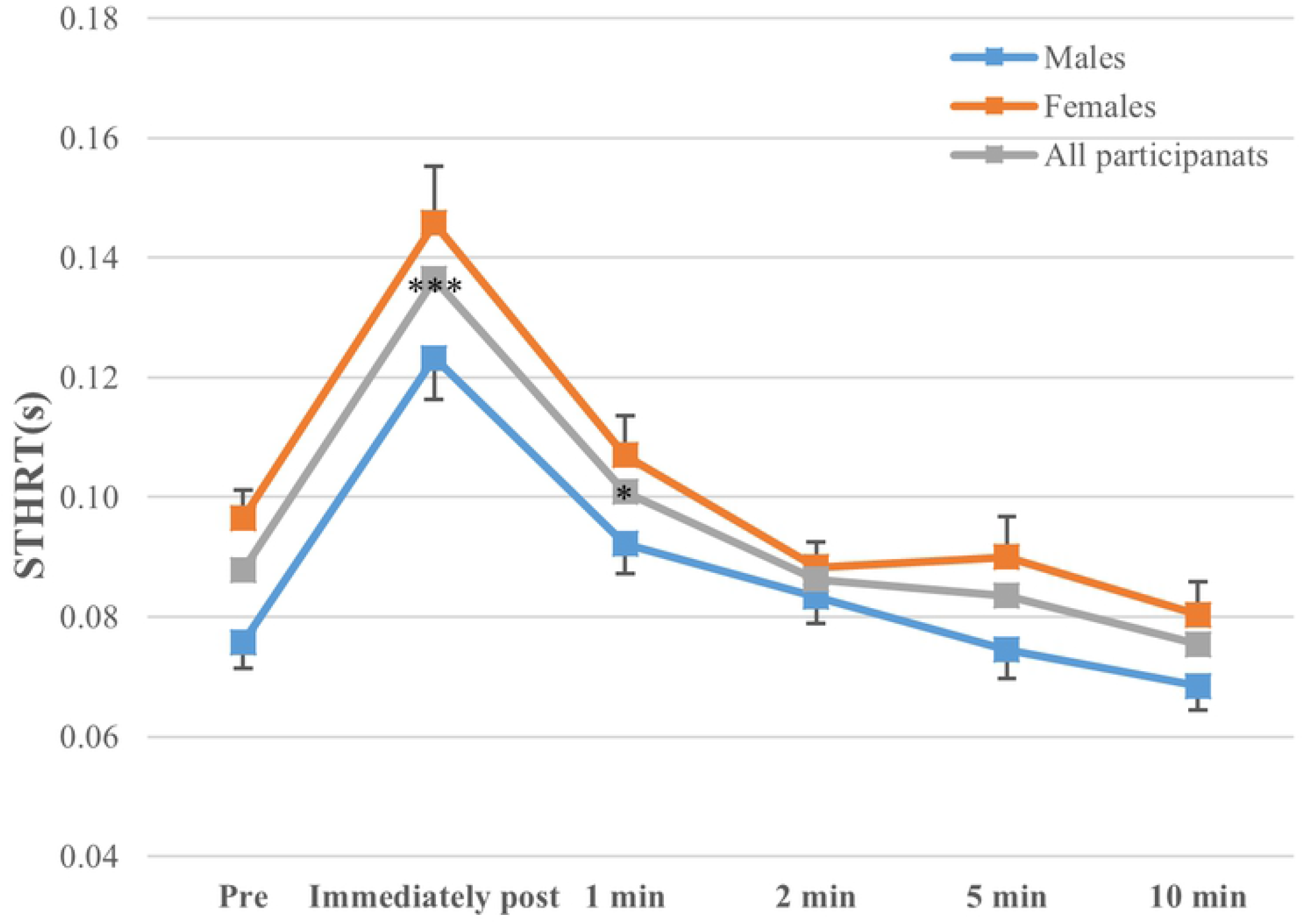
Changes in singlet twitch half-relaxation time (STHRT) with fatigue and recovery. *** (*p*<0.001) and * (*p*<0.05) indicates a significant increase in the relaxation time at post-exercise compared to pre-exercise values.

**Fig 7.**
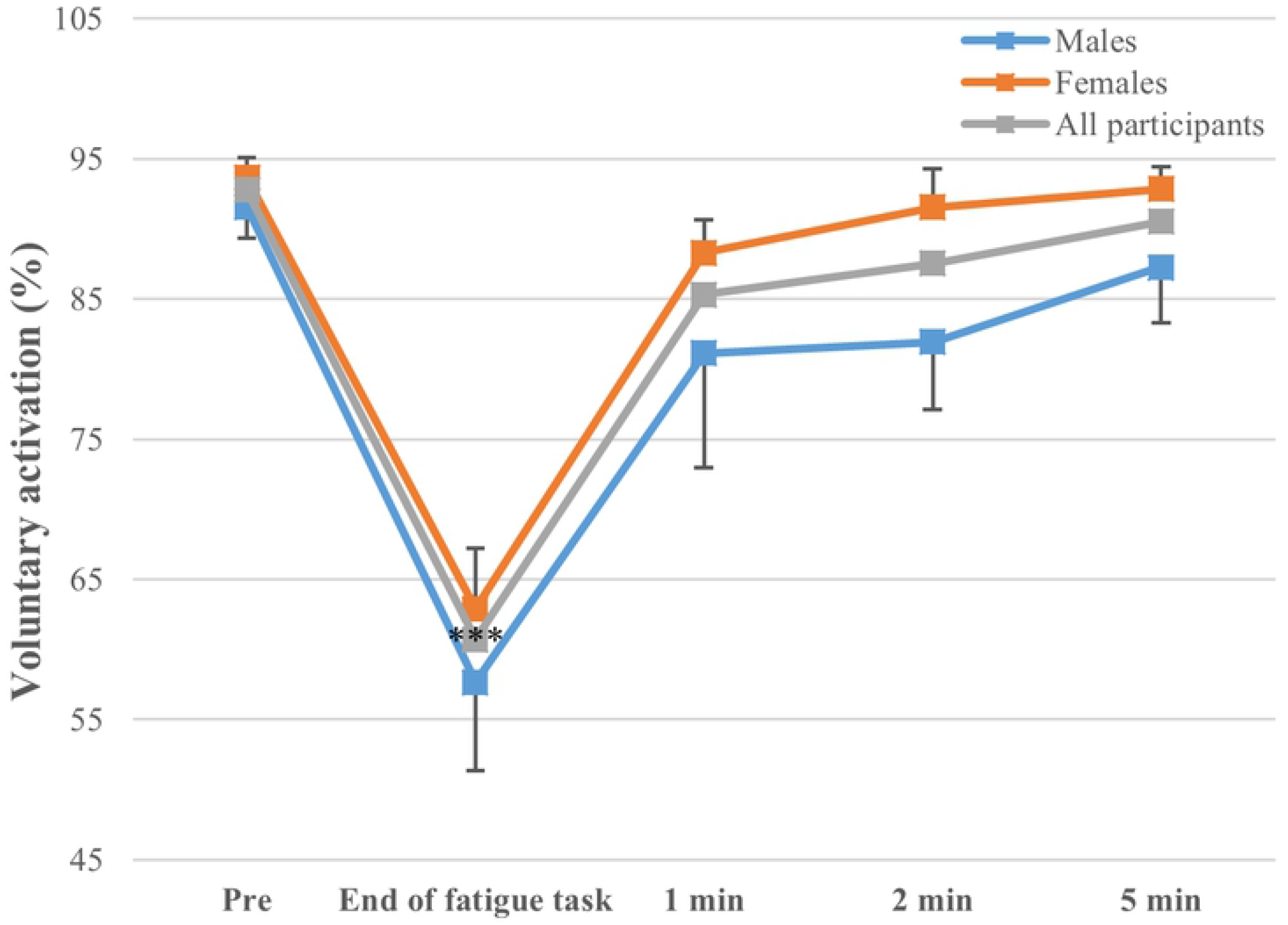
Changes in voluntary activation with fatigue and recovery. *** (*p*<0.001) indicates a significant decrease in the voluntary activation at post-exercise compared to pre-exercise values (Pre).

**Fig 8.**
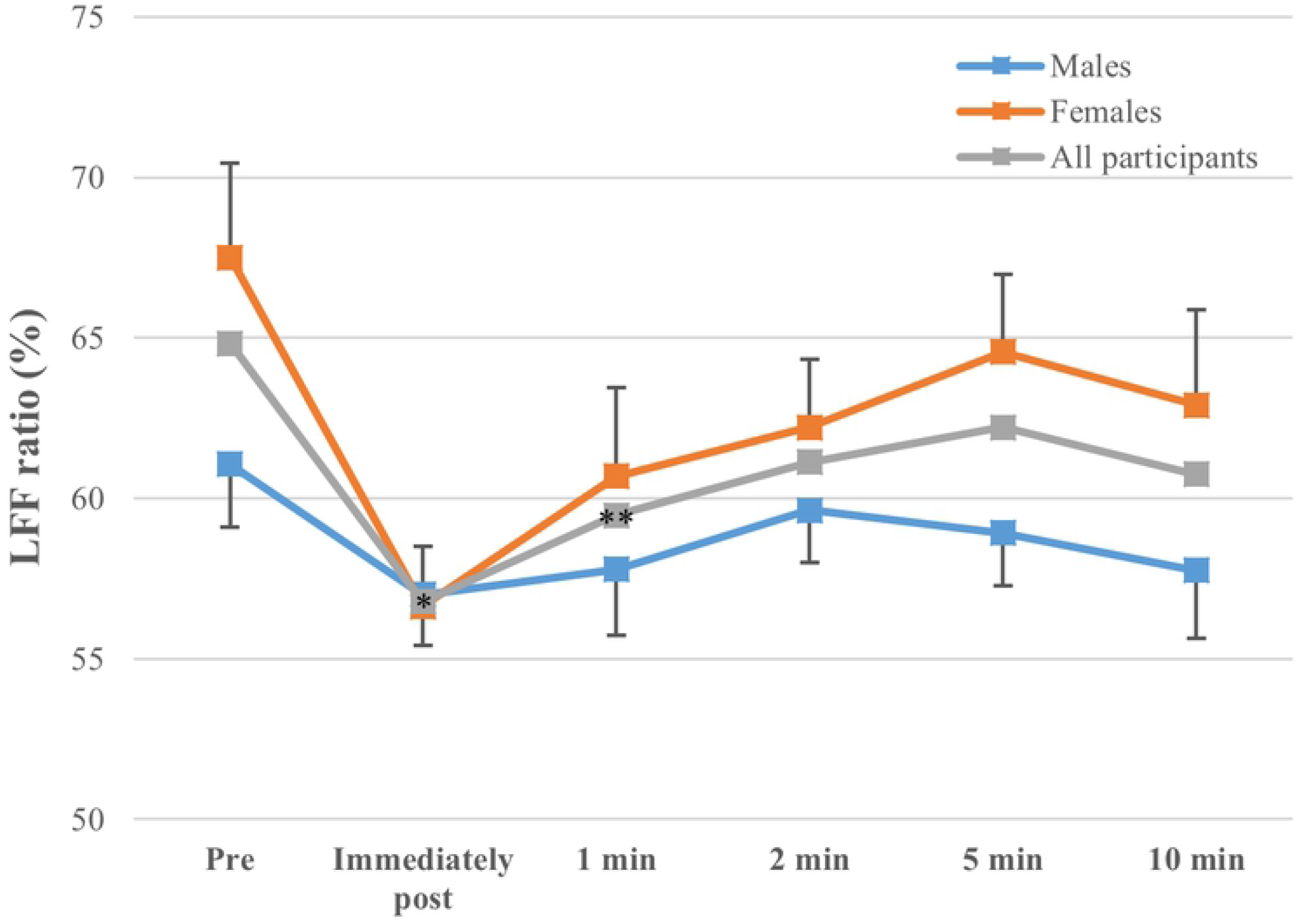
Changes in Low frequency fatigue (LFF) ratio with fatigue and recovery. * (*p*<0.01) and ** (*p*<0.01) indicate a significant reduction in the low frequency fatigue ratio from pre to post-exercise.

### Fatigue effects and interactions

Fatigue had a significant effect on all dependent variables, and, affected men and women similarly (no sex X fatigue interaction), except for MVIC torque (Table 1) showing a somewhat greater decline with fatigue in males (∼28 %) compared with females (∼22 %) (Fig 2). MVIC torque was decreased after fatigue and not fully recovered at 10 min post-fatigue (Fig 2). Measures reflecting changes in central (voluntary activation) and peripheral mechanisms (twitch torque-related variables) were all affected by fatigue, but all recovered within 5min post-fatigue (Figs 3-8).

The results of the Pearson correlation and multiple regression analyses performed in an attempt to explain the change in MVIC torque 1 min after the end of the fatigue task are shown in Tables 2 and 3, respectively. Both changes in voluntary activation and twitch torque (doublet and singlet) were positively correlated with the change in MVIC torque. Furthermore, 71% of the variance in MVIC torque decrease after 1 min of recovery could be explained by the multiple linear regression model, in which changes in voluntary activation and singlet twitch peak torque were identified as significant influencing factors.

**Table 2.** Pearson’s correlations between pre-to-post (1min) changes in MVIC torque and central/peripheral fatigue properties.

| Items | r | 95% CI of r | p-values |
| --- | --- | --- | --- |
| MVIC and DT | 0.380 | 0.027–0.709 | 0.067 |
| MVIC and ST | **0.419\*** | -0.047–0.774 | **0.042** |
| MVIC and VA | **0.755\*\*** | 0.406–0.891 | **<0.001** |

\*\* = $p < 0.01$ ; \* = $p < 0.05$ ; MVIC: maximal voluntary isometric contraction; DT: resting doublet twitch peak torque; ST: resting singlet twitch peak torque; VA: voluntary activation.

**Table 3.**
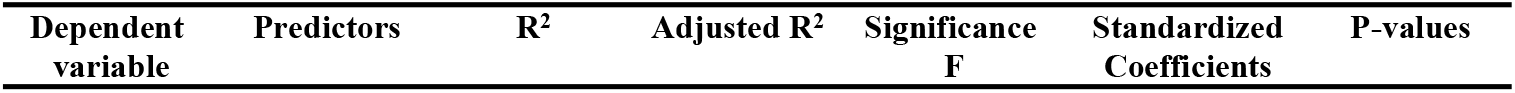
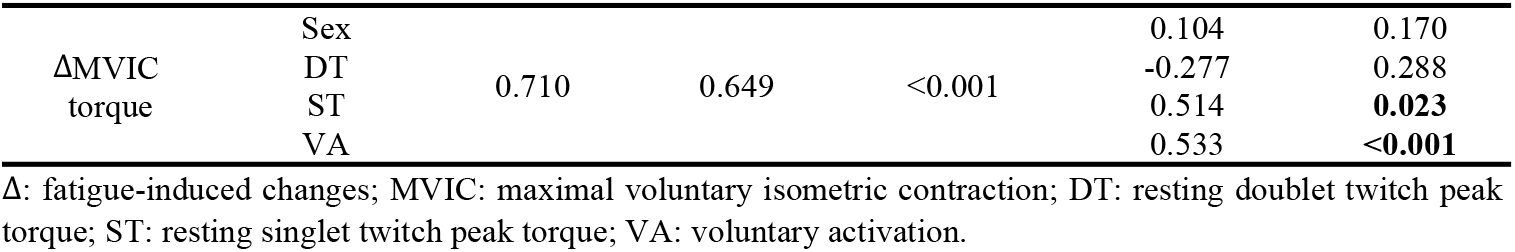
Multiple linear regression model with regards to the contribution of peripheral and central fatigue parameters (predictors) to fatigue-induced changes (pre to 1min post-exercise) in MVIC torque.

## Discussion

The main aim of this study was to determine sex differences in central and peripheral fatigue produced by isometric ankle plantar flexors exercise in healthy young adults. There was a significant effect of fatigue on all variables suggesting the presence of both central and peripheral fatigue, with no significant sex differences in the observed fatigue-related changes (other than MVIC torque).

### Overall fatigue and sex differences

We observed a significant exercise-induced decrease in MVIC peak torque, which had not fully recovered 10 min following exercise completion. This is in line with previous findings. The 24% decline in MVIC torque from before to 1 min after exercise in our study is in line with a 30% reduction observed immediately after a 25% MVIC isometric exercise performed with ankle plantar flexors until failure (10). The prolonged reduction in MVIC torque up to 10 min post-exercise has also been shown in studies of ankle plantarflexors (29) and other muscle groups such as elbow flexors, with reductions still shown at 10 min (21) and even 45 min (23) following sustained isometric exercise in healthy young adults.

Interestingly, males presented with a greater decrease (∼28%) in MVIC peak torque from pre-fatigue to post-fatigue (average of the recovery period) compared with females (∼22%), possibly indicating greater fatigability in males, as found in several previous studies using sustained isometric exercise of various muscle groups, including ankle dorsiflexors (14,20–23). This sex difference is also acknowledged in review studies (12,18), which conclude that although sex differences in fatigability depend on task specific features including intensity, exercising muscles and contraction type/speed, males are usually more fatigable than females for isometric fatiguing contractions performed at a similar relative intensity. However, our results indicate that the difference in fatigability of ankle plantar flexor muscles between males and females is relatively small in comparison to other muscle groups (12). In further support of this, fatiguing exercise duration was not statistically different between males and females, although the females tended to exercise longer before reaching the set fatigue criteria (by about 40 s on average, a 17% difference between sexes).

The greater fatigability in males compared with females has been explained by different potential factors, which can be inter-related and include (in males): greater muscle mass and strength, greater blood flow occlusion leading to a greater accumulation of metabolites and lesser muscle perfusion, a lesser proportion of fatigue-resistant muscle fibers (type I) within skeletal muscles and a greater failure in voluntary activation (12,18). Concerning the small but significant difference reported here, it does not appear to be related to strength, as there was no association between MVIC torque and time to fatigue (r = -0.15; p = 0.48). Because ankle plantar flexor muscles can present with lesser differences in relative area of type I muscle fibers between males and females compared with other muscle groups (30–33), this could explain, at least in part, the smaller difference in fatigability between the two groups reported in the present study. In contrast with other lower extremity muscle groups (14,19), no sex difference in central fatigue were observed in the present study, thus eliminating this factor as a major contributor to the somewhat greater fatigability of males we report.

### Central fatigue

We found a significant decrease in voluntary activation at the end of the fatigue task, when participants were assumed to be producing a maximal effort but failing to reach the 30% of MVIC target (torque). Even though voluntary activation was approximately 7 % and 5 % lower than before fatigue after 1 and 2 min of recovery, post hoc tests failed to detect significant differences between pre-fatigue values and any values during the recovery period. These results are partly in line with previous findings showing reduced voluntary activation of ankle plantar flexor muscles immediately after a sustained 25% MVIC until exhaustion (10) and up to at least 2 min post-exercise involving intermittent bilateral MVICs until torque decreased by 50% (29). In the present study, voluntary activation was recovered (statistically) 1 min after the termination of the fatigue task. A rapid recovery of the voluntary activation has been shown in previous studies (e.g. Bilodeau 2006 (34)). Our findings did not show a significant sex difference in central fatigue, even though males had a somewhat more pronounced decrease in voluntary activation and slower recovery compared with females (Fig 7). This lack of a significant difference between sexes could be explained by the greater variability in voluntary activation observed in post-fatigue of males. However, in contrast to what has been reported for other lower extremity muscle groups (e.g., leg extensors and ankle dorsiflexors) (14,19), ankle plantarflexor muscles may not exhibit an important sex difference with regards to the extent of central fatigue, at least for a relatively low-level sustained isometric fatigue task. Interestingly, the results of the multiple regression analysis (Tables 3) suggest that central fatigue was as important as peripheral fatigue in explaining the decreased MVIC torque at 1 min post-fatigue. Thus, it appears that the somewhat greater fatigability of males shown here could involve both central and peripheral factors, even though no significant sex X fatigue interactions were found when those factors are taken in isolation (see also next section).

### Peripheral fatigue

Our results showed fatigue effects on twitch contractile properties, as reported in previous studies: reduced singlet/doublet twitch peak torques (8,20,22), decreased low frequency fatigue ratio (11,35), and increased singlet and doublet twitch half-relaxation time (9,10,19,36). These changes reflect potential alterations in neurotransmission and/or excitation–contraction coupling mechanisms and reflect the presence of peripheral fatigue (1,37). More specifically, the 19% reduction in doublet twitch peak torque immediately post-exercise observed here (Fig 3) is in accordance with prior results from our group (10) using a similar fatigue criterion (25% MVIC) in the same muscle group, with a 10% fatigue-induced reduction in doublet twitch peak torque. A sustained reduction in singlet twitch torque up to 10 min post-exercise is also in line with previous findings (14) showing a 14-min sustained decrease in singlet twitch force following exercise of ankle dorsiflexors. Such decreases in twitch peak torque potentially reflect metabolic inhibition of the muscle contractile processes, such as resulting from the fall in pH level within the fatigued muscle (38). Also, a decrease in the low frequency fatigue ratio was statistically noted by 1min after the fatigue task (Fig 8). This reduction potentially indicates decreased Ca ^2+^ release from the sarcoplasmic reticulum and a related decrease in the amount of Ca ^2+^ binding to myofilaments possibly limiting the amount of muscle force produced (11,35).

Significant increases in doublet and singlet half-relaxation time were shown by 1 min after exercise (Figs 5 and 6). Such slowing of muscle relaxation may result from slowing of Ca ^2+^ re-uptake by the sarcoplasmic reticulum and slowed Ca^2+^ detachment from myofilaments (39,40). Doublet and singlet half-relaxation times were also found to be longer in females than males. This is consistent with previous literature (19,20,36) and has been explained by a greater type I muscle fiber area in females or by differences in other mechanisms involved in muscle relaxation, including Ca^2+^ re-uptake and Ca^2+^ detachment mentioned above. However, such difference was not accompanied by different changes in half-relaxation time with fatigue between males and females in the present study. Again, it is possible that the difference in relative area of type I muscle fibers between males and females in ankle plantar flexors compared with other muscle groups (30–33) may not be important enough to lead to a sex difference in fatigue-related changes in twitch half-relaxation time.

## Conclusion

This study aimed to determine potential differences between males and females in central and peripheral fatigue of ankle plantar flexor muscles in healthy young adults. Altogether, our findings showed no major differences in central or peripheral fatigue between males and females to explain a somewhat greater fatigability in males.

## Supporting information

**S1 file. Data set**. This data includes maximal voluntary isometric contraction voluntary (MVIC) torque and central/peripheral fatigue variables which were collected before and immediately after exercise (or at the end of exercise), and recovery period (1, 2, 5 10 min), as well as the time to fatigue.

## Author contributions

**Conceptualization:** Donguk Jo, Miriam Goubran, Martin Bilodeau.

**Data curation:** Donguk Jo, Miriam Goubran.

**Formal analysis:** Donguk Jo, Martin Bilodeau

**Funding Acquisition:** Martin Bilodeau

**Investigation:** Donguk Jo, Miriam Goubran, Martin Bilodeau.

**Methodology:** Donguk Jo, Miriam Goubran, Martin Bilodeau.

**Project administration:** Donguk Jo, Miriam Goubran, Martin Bilodeau.

**Resources:** Martin Bilodeau

**Software:** Donguk Jo, Martin Bilodeau

**Validation:** Donguk Jo, Miriam Goubran, Martin Bilodeau

**Visualization:** Donguk Jo

**Writing (original draft):** Donguk Jo, Martin Bilodeau.

Writing (review/edits): Donguk Jo, Miriam Goubran, Martin Bilodeau.

## Notes

### Competing Interest Statement

The authors have declared no competing interest.

